# Mutations in the tomato gibberellin receptors suppresses xylem proliferation and reduces water loss under water-deficit conditions

**DOI:** 10.1101/2020.01.06.895011

**Authors:** Natanella Illouz-Eliaz, Idan Nissan, Ido Nir, Uria Ramon, Hagai Shohat, David Weiss

## Abstract

Low gibberellin (GA) activity in tomato (*Solanum lycopersicum*) inhibits leaf expansion and reduces stomatal conductance. These lead to lower transpiration and improve water status under transient drought conditions. Tomato has three GIBBERELLIN-INSENSITIVE DWARF1 (GID1) GA receptors with overlapping activities and high redundancy. We have tested whether mutation in a single GID1 reduces transpiration without affecting growth and productivity. CRISPR-Cas9 *gid1* mutants were able to maintain higher leaf water content under water-deficit conditions. Moreover, while *gid1a* exhibited normal growth, it showed reduced whole plant transpiration and better recovery from dehydration. Mutation in *GID1a* inhibited xylem vessels proliferation that led to lower hydraulic conductance. In stronger GA mutants, we also found reduced xylem vessel expansion. These results suggest that low GA activity affects transpiration by multiple mechanisms; it reduces leaf area, promotes stomatal closure and reduces xylem proliferation and expansion and as a result, xylem hydraulic conductance. We further examined if *gid1a* perform better than the control M82 in the field. Under these conditions, the high redundancy of GID1s was lost and *gid1a* plants were semi-dwarf, but their productivity was not affected. Although *gid1a* did not perform better under drought conditions in the field, it exhibited higher harvest index.

**Highlight:** The loss of the tomato gibberellin receptors GID1s reduced xylem proliferation and xylem hydraulic conductance. These contribute to the effect of low gibberellin activity on water loss under water-deficit condition.

## Introduction

Drought has a major impact on plant development and food supply, and is responsible for major losses of crop productivity (Mittler and Blumwald, 2010). Plants have adopted various strategies to cope with water deficiency, including maintaining water status by stomatal closure, accumulation of osmolytes and stress related proteins and changes in growth and development (Skirycz and Inzé, 2010; Osakabe et al., 2014). Rapid stomatal closure, expression of stress related genes and developmental changes in response to water deficiency are mediated primarily by the stress hormone abscisic acid (ABA; Cutler *et al*., 2010). Several studies suggested that the ABA-antagonist hormone, gibberellin (GA), has also role in these responses (Colebrook *et al*. 2014).

GA regulates numerous developmental processes throughout the life cycle of the plant, from germination to fruit development (Davière and Achard, 2013). All GA responses are suppressed by the nuclear DELLA proteins (Locascio *et al*., 2013; Livne *et al*., 2015). GA binding to its receptor GIBBERELLIN-INSENSITIVE DWARF1 (GID1) increases the affinity of the latter to DELLA. The formation of the GID1-GA-DELLA complex recruits an F-Box protein SLEEPY1 (SLY1) to DELLA, leading to DELLA polyubiquitination and degradation in the proteasome (Harberd *et al*., 2009). This initiates transcriptional reprograming and activation of GA responses (Hauvermale *et al*., 2012).

GID1 was first discovered in rice, and the rice mutant *gid1-1* is extremely dwarf and insensitive to GA (Ueguchi-Tanaka *et al*., 2005). While rice, similarly to other monocots, has a single GID1 gene, Arabidopsis has three homologues with partially overlapping functions (Griffiths *et al*., 2006; Nakajima *et al*., 2006). Similarly, tomato (*Solanum lycopersicum*) has three GA receptors; GID1a, GID1b1 and GID1b2. These receptors exhibit high redundancy under optimal controlled growth conditions, but under extreme ambient conditions, all three are required for robust growth (Illouz-Eliaz *et al*., 2019). While *gid1b1* and *gid1b2* single mutants do not show clear phenotype, *gid1a* is slightly shorter with darker green leaves. GID1a is the dominant GA receptor in the regulation of germination, stem elongation and leaf expansion and exhibits the highest affinity to the single tomato DELLA protein PROCERA (PRO, Illouz-Eliaz *et al*., 2019).

Recent studies have shown that altering GA levels or signal, improve plant tolerance to water-deficit stress (Colebrook *et al*., 2014). Inhibition of GA biosynthesis by paclobutrazol (PAC) increased tolerance to water deficiency in cereals (Plaza-Wuthrich *et al*., 2016) and tomato (Pal *et al*., 2016). Ectopic expression of *MhGAI1* (the tea crabapple DELLA gene) in tomato, promotes drought tolerance (Wang *et al*. 2011). Inhibition of GA activity in tomato by overexpressing the Arabidopsis *GA METHYL TRANSFERASE 1* (*AtGAMT1*) gene or the gain-of-function stable DELLA mutant gene *proΔ17*, reduced whole plant transpiration and improved resistance to drought (Nir *et al*., 2014; Nir *et al*., 2017). Several possible mechanisms for this stress tolerance were suggested, including indirect effects on transpiration due to reduced plant size (Magome *et al*., 2008; Achard *et al*., 2006) and direct effect on transpiration, due to increased response to ABA in guard cell and rapid stomatal closure (Nir et al., 2017). Low GA activity also led to the activation of various stress-related genes (Wang *et al*., 2008; Tuna *et al*., 2008) and the accumulation of osmolytes (Omena-Garcia et al., 2019). GA also affects vascular development; it promotes xylem expansion and secondary vascular development (Ragni et al, 2011; Dayan et al., 2012; Aloni, 2013). Xylem vessel area can affect hydraulic conductance and water status in response to environmental changes (Melcher et al, 2012; Brodribb, 2009).

GA has a pleotropic effect on plant development. Since the three tomato GA receptors exhibit high redundancy in the regulation of growth, we examined here if mutation in a single GID1 can improve drought tolerance without affecting growth and productivity. Our results show that mild attenuation of GA activity due to the loss of *GID1a* was sufficient to reduce whole plant transpiration and water loss under water-deficit conditions without affecting plant growth. They also suggest that low GA activity affects transpiration by multiple mechanisms; it inhibits leaf growth, promotes stomatal closure, and reduced xylem vessels proliferation and expansion and therefore hydraulic conductivity.

## Materials and methods

### Plant materials and growth conditions

Tomato cv M82 (sp^-^/sp^-^) plants were used throughout this study. The CRISPR-Cas9 *gid1* and *sly1* mutants (Illouz-Eliaz et al., 2019) were in the M82 background. Plants were grown in a growth room set to a photoperiod of 12/12-h night/days, light intensity) cool-white bulbs) of ∼250 μmol m^-2^ s^-1^, and 25°C. In other experiments, plants were grown in a greenhouse under natural day-length conditions, light intensity of 700 to 1000 µmol m^-2^ s^-1^ and 18-30°C. In the summer (April to August) of 2019 *gid1* single mutant lines and M82 were grown in an open field under ambient conditions (Acre, Israel).

### Tomato *SLY1* CRISPR/Cas9 mutagenesis, plant transformation and selection of mutant alleles

Two single-guide RNAs (sgRNAs, Supplemental Table 1) were designed using the CRISPR-P tool (http://cbi.hzau.edu.cn/crispr). Vectors were assembled using the Golden Gate cloning system as described in Weber et al. (2011). Final binary vector, pAGM4723, was introduced into *Agrobacterium tumefaciens* strain GV3101 by electroporation. The construct was transferred into M82 cotyledons using transformation and regeneration methods described by McCormick (McCormick, 1991). Kanamycin-resistant T0 plants were grown and transgenic lines were selected and self-pollinated to generate homozygous transgenic lines. For genotyping of the transgenic lines, genomic DNA was extracted, and each plant was genotyped by PCR for the presence of the Cas9 construct. The CRISPR/Cas9-positive lines were further genotyped for mutations in *SlSLY* (Solyc04g078390) using a forward primer to the left of the sgRNA1 target sequence and a reverse primer to the right of the sgRNA2 target sequence.

### Relative water content (RWC) determination

Leaf RWC was measured as follows: fresh leaf weight (FW) was measured immediately after leaf detachment. Leaves were then soaked for 8 h in 5 mM CaCl_2_ in the dark at room temperature, and the turgid weight (TW) was recorded. Dry weight (DW) was recorded after drying the leaves at 70°C for 48 h. RWC was calculated as (FW− DW)/(TW − DW) x 100 (Sade *et al*., 2009).

### Measurements of stomatal index and density

Stomatal index (stomatal number/total number of epidermal cells) and stomatal density were determined using the rapid imprinting technique (Geisler *et al*., 2000). This approach allowed us to reliably and simultaneously score hundreds of stomata from each experiment. Briefly, vinylpolysiloxane dental resin (eliteHD+; Zhermack Clinical) was attached to the abaxial side of the leaf, dried for 1 min, and then removed. The resin epidermal imprints were covered with transparent nail polish, which was removed once it dried and served as a mirror image of the resin imprint. The nail polish imprints were placed on glass cover slips and photographed under a model 1M7100 bright-field inverted microscope (Zeiss, Jena, Germany) with a mounted Hitachi HV-D30 CCD camera (Japan).

### Measurement of Leaf Area

Total leaf area was measured in six weeks-old M82, *gid1a* and *gid1a gid1b2* plants, using a model Li 3100 leaf area meter (LI-COR Biosciences, Lincoln, NE, USA).

### Whole-plant transpiration, transpiration rate and whole canopy conductance measurements

Whole-plant transpiration ratewas determined using an array of lysimeters placed in the greenhouse (Plantarry 3.0 system; Plant-DiTech) in the “iCORE Center for Functional Phenotyping” (http://departments.agri.huji.ac.il/plantscience/icore.phpon), as described in detail by Halperin *et al*. (2017). Briefly, plants were grown in 4L pots under semi-controlled temperature conditions (20–32°C night/day), natural day-length, and light intensity of approximately 1000 µmol m^-2^ s^-1^. Each pot was placed on a temperature-compensated load cell with digital output (Vishay Tedea-Huntleigh) and sealed to prevent evaporation from the surface of the growth medium. The weight output of the load cells was monitored every 3 min. The data were analyzed using SPACanalytics (Plant-Ditech) software to obtain the following whole-plant physiological traits: Daily plant transpiration (weight loss between predawn and sunset) was calculated from the weight difference between the two data points. Whole canopy conductance (G*sc*) was calculated by dividing *E* (transpiration rate/plant weight) by vapor pressure deficit (VPD). The plant daily weight gain (ΔPWn) between consecutive days was:

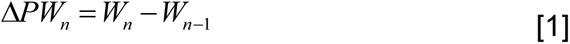

where Wn and Wn-1 are the container weights upon drainage termination on consecutive days, n and n-1. Following Eq. 2, the weight on day n is the sum of plant weight on day n-1 and the weight gain ΔPWn-1

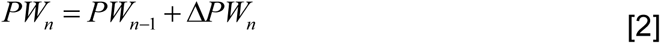

The whole-plant WUE during a defined period was determined by the ratio between the sum of the daily plant fresh-weight gain (ΔPW) and water consumed throughout this period (comulitative daily transpiration -PDT):

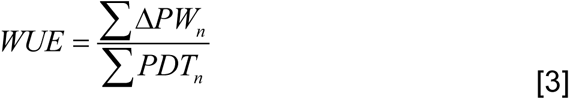

### Isolation of guard cells for qRT-PCR analysis

Guard cells from tomato leaves were isolated according to Nir *et al*. (2017). Briefly, 20 g of fully expanded leaves without the veins were ground twice in a blender in 100ml cold distilled water, each time for 1 min. The blended mixture was poured onto a 200-µm mesh (Sefar AG, Heiden, Switzerland) to remove mesophyll and broken epidermal cells. The remaining epidermal peels were rinsed thoroughly with deionized water. The peels were then transferred into 10-ml buffer (Araújo *et al*., 2011) containing the enzyme CELLULYSIN cellulase from *Trichoderma viride* (Calbiochem, La Jolla, CA, USA) and digested for 1 h at a shaking speed of 150 rpm. This enzymatic treatment digests pavement cells, but not guard cells (Wang *et al*., 2011). The digested material was poured again onto a 200-µm mesh placed in a tube and rinsed thoroughly with digestion buffer (without the enzyme). To remove residues of buffer and cell particles, the tubes were centrifuged at 4°C for 5 min at 2200 rpm. Samples of digested epidermal strips were stained with neutral red, and cell vitality was examined microscopically (Nir *et al*., 2017).

### qRT-PCR analysis

qRT-PCR analysis was performed using an Absolute Blue qPCR SYBR Green ROX Mix (AB-4162/B) kit (Thermo Fisher Scientific, Waltham, MA USA). Reactions were performed using a Rotor-Gene 6000 cycler (Corbett Research, Sydney, Australia). A standard curve was obtained using dilutions of the cDNA sample. The expression was quantified using Corbett Research Rotor-Gene software. Three independent technical repeats were performed for each sample. Relative expression was calculated by dividing the expression level of the examined gene by that of *ACTIN*. Primer sequences are presented in Supplemental Table 1.

### Measurements of hydraulic conductance

Measurements of volumetric flow-rate, to determine hydraulic conductance, were performed according to Melcher *et al*. (2012) with some modifications. Three cm long segments were dissected from the stems from the same location. The top of the segments were connected via silicone tubing to pipet containing 15mM KCl, and mounted vertically, while the bottom end of the stem was connected to a drainage tube. To calculate the hydraulic conductance (K’) we have used the following equation:

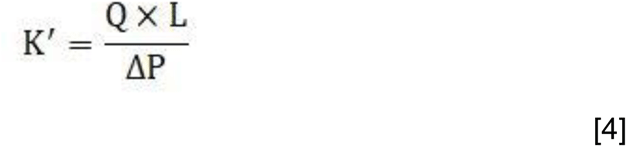

The volume of fluid, which passed through the stem during a constant time interval was measured to calculate the volumetric flow rate (Q; mmol H_2_O/sec), L-length of the stem segment (m), ΔP – pressure (the driving force, MPa, calculated by the hydraulic-head height). All dissections and connection of the apparatus were performed under water to avoid embolism.

### Microscopic analysis of the xylem

Stem or petiole segments were manually dissected to thin cross-section slices using a razor blade. The cross sections were then stained using a modified Weisner reaction (Pradhan Mitra and Loqué, 2014), which stains the lignin in the xylem vessels. The stained cross sections were examined under a LEICA ICC50W light microscope. The images were then manually analyzed, using ImageJ software (http://rsb.info.nih.gov/ij/), xylem vessel area, diameter and number were measured.

### Calculation of theoretical specific hydraulic conductivity (Kts)

To evaluate xylem specific hydraulic conductivity, we used the modified Hagen-Poiseuille equation (Tyree and Ewers, 1991) which calculates the theoretical hydraulic conductivity (Kt; mmol m MPa-1 s-1) of a bundle assuming perfectly cylindrical pipes:

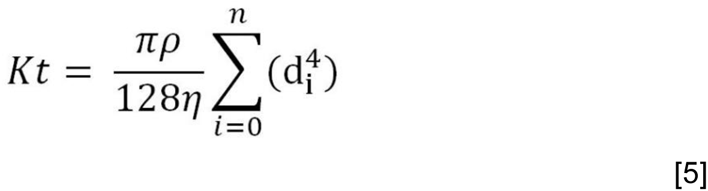

Where d is the vessels diameter, ρ is the fluid density in kg x m^-3^, η is the fluids dynamic viscosity in MPa s^-1^, and n is the number of pipes in the bundle. The theoretical specific hydraulic conductivity (Kts; mmol m-1 s-1 MPa-1) was calculated by normalizing Kt to leaf area (LA) (Hochberg *et al*., 2015):

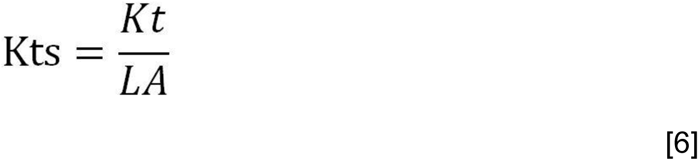

Leaf area was calculated by scanning the foliage (LaserJet pro 400 MFP M475dw), and measuring the leaf area with ImageJ (http://rsb.info.nih.gov/ij/).

## Results

### The loss of *GID1* reduced water loss and whole plant transpiration under water-deficit conditions

To examine the contribution of the three GID1 receptors to plant water status, we first compared the rate of water loss in M82 and all single and double *gid1* mutants under water-deficit conditions. All plants were grown until they produced five expanded leaves, after which irrigation was stopped and the soil was allowed to dry out progressively. After 7 days, non-irrigated M82, *gid1b1*, *gid1b2* and the double mutant *gid1b1 gid1b2* plants began to wilt, whereas *gid1a*, *gid1a gid1b1* and *gid1a gid1b2* lines remained turgid. At this time point, we measured relative water content (RWC) of the leaves. RWC in M82, *gid1b1*, *gid1b2* and *gid1b1 gid1b2* was reduced (compared to irrigated plants) by approximately 20%, while in *gid1a*, *gid1a gid1b1* and *gid1a gid1b2* RWC was similar to the irrigated plants (Fig. 1A). We continued the drought treatment to M82 and *gid1a gid1b2* plants and after three more days, *gid1a gid1b2* plants also wilted. Four days later, plants were rehydrated and their ability to recover was monitored. M82 plants failed to recover, but *gid1a gid1b2* plants fully recovered and necrotic lesions were found only on several leaves (Fig. 1B). These results suggest that the loss of GID1, similar to increased DELLA activity (Nir et al., 2017), reduces water loss under water deficit conditions. They also propose that GID1a has the most prominent role in this process.

**Fig. 1.**
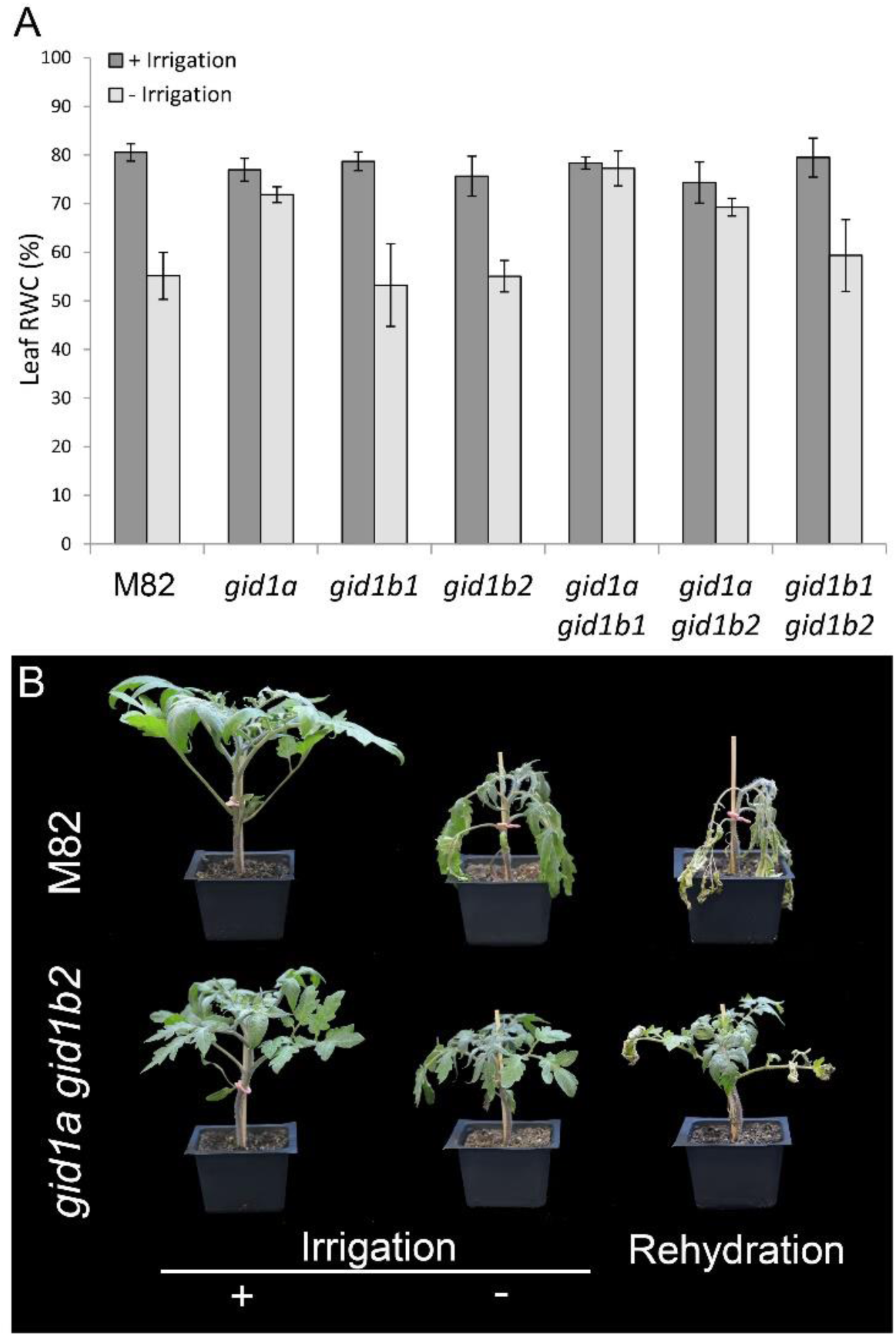
The *gid1*s exhibit reduced water loss under water-deficit conditions. **A.** Average leaf relative water content (RWC) of control M82 and *gid1* single and double mutants grown with or without irrigation for 7 days ± SE. Values are means of eight replicates ± SE. **B.** Representative M82 and *gid1a gid1b2* plants grown under normal irrigation regime (+irrigation) or without irrigation for 7 days. After 14 days without irrigation, plants were rehydrated and recovery was assessed after 10 days.

Leaf area in four-weeks-old *gid1a gid1b2* was smaller, but in *gid1a*, similar to M82 (Fig. 2A). Since *gid1a* exhibited reduced water loss but similar leaf area to M82, we further focus on this line. We first analyzed microscopically the abaxial leaf epidermal tissues of M82 and *gid1a.* We did not find significant differences in stomatal index (Supplementary Fig. S1A), suggesting that the loss of *gid1a* does not change the ratio between pavement cells and guard cells. Also stomatal density was not affected by the mutation (Supplementary Fig. S1B), suggesting that the total number of stomata in *gid1a* is similar to M82. Previously we showed that all double mutants exhibit reduced whole plant transpiration (Illouz-Eliaz et al., 2019). Here we examined whole-plant transpiration in irrigated M82 and *gid1a* mutant plants grown in a greenhouse using an array of load cells (lysimeters) which simultaneously followed the daily weight loss of each plant (Nir et al., 2017). Daily transpiration, transpiration rate and whole canopy conductance of *gid1a* were significantly lower than that of M82 (Fig. 2B,C,D). Since transpiration of *gid1a* was lower than M82 but their growth was similar, the water use efficiency (WUE) of *gid1a* was higher than that of M82 (Fig. 2E). These results imply that mutations in GA receptors promote stomatal closure similar to the effect of stable DELLA overexpression (*35S:proΔ17*, Nir et al., 2017). We therefore tested if the three GID1s are expressed in guard cells. To this end, we isolated guard cells from M82 and analyzed the expression of the three genes in guard-cell enriched samples. All GID1 genes were expressed in guard cells and *GID1a* exhibited the highest expression (Fig. 2F).

**Fig. 2.**
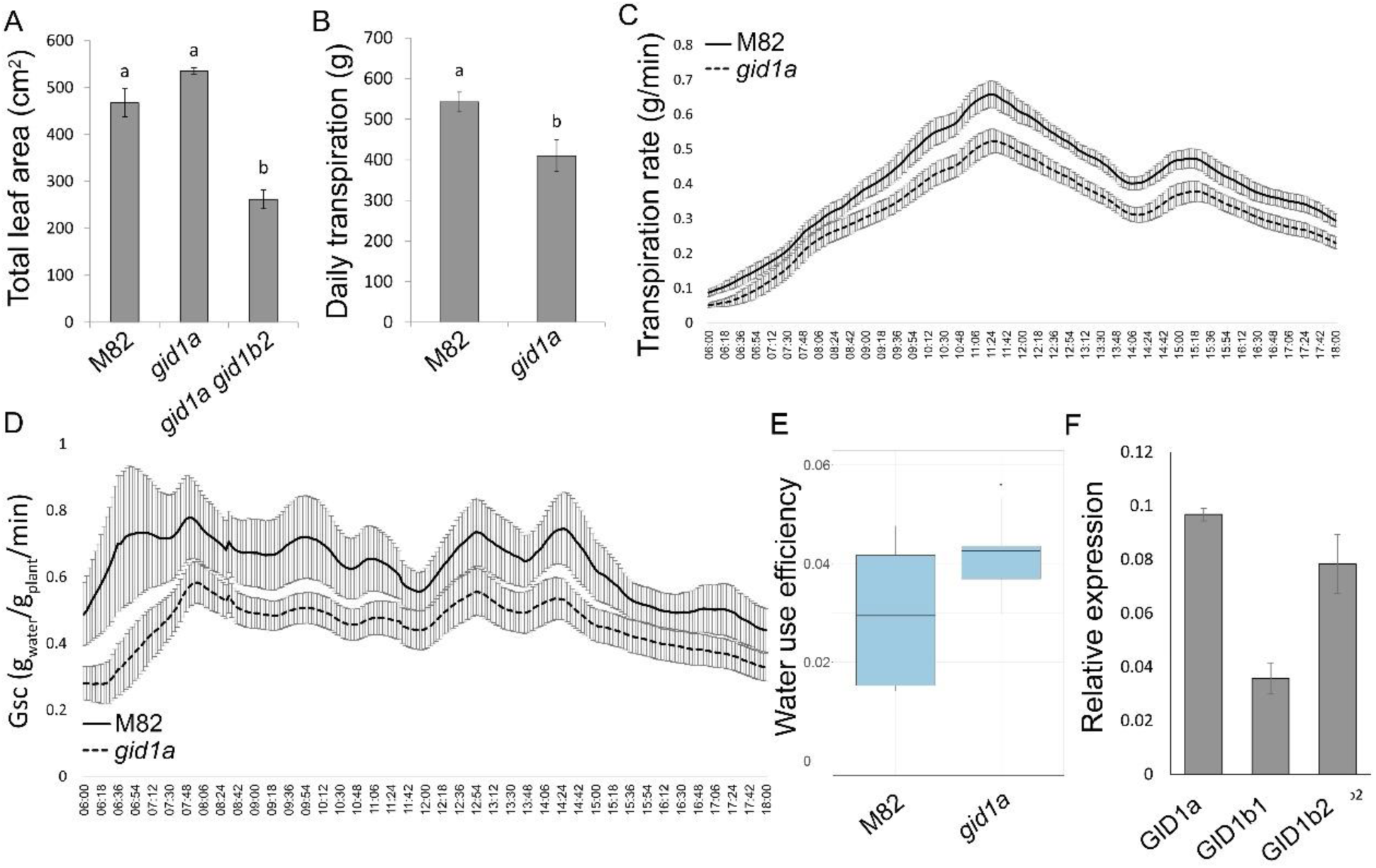
Loss of GID1a reduced whole plant transpiration. **A.** Total leaf area of control M82, *gid1a* and *gid1a gid1b2* six-weeks-old plants. Values are mean of 9 plants ± SE. Small letters represent significant differences between the lines (Student’s t test, P<0.05). **B.** Whole plant daily transpiration of M82 and *gid1a*. Plants were placed on lysimeters and pot (pot + soil + plant) weight was measured every 3 min. Values are means of 13 plants ± SE. Each set of letters above the columns represents significant differences between respective treatments (Student’s t test, P<0.05). **C.** Whole-plant transpiration rate over the course of 12 h (06:00 AM to 06:00 PM). Values are means of 13 plants ± SE. **D.** Whole canopy conductance (Gsc) of M82 and *gid1a* (calculated by dividing *E* (transpiration rate/plant weight) by vapor pressure deficit (VPD). Values are means of 13 plants ± SE.. **E**. Whole plant water use efficiency (WUE) of M82 and *gid1a* was calculated as the ratio between plant growth and transpirationData (taken from 13 different plants) are graphically presented as whisker and box plots. **F**. qRT-PCR analysis of *GID1* expression in M82 isolated guard cells. Values are means of three biological replicates ± SE.

Next we tested the effect of water-deficit conditions on transpiration rate in M82 and *gid1a* plants. After two weeks of growth on the lysimeters, we gradually reduced irrigation (each day by 50%) to expose M82 and *gid1a* plants to water-deficit conditions. In the first four days of the water-deficit treatment, transpiration rate in M82 was higher than in *gid1a* (Fig. 3A). However, at day 6, as water availability in the pots become a limiting factor, transpiration rate of M82 rapidly declined. On the other hand, transpiration of *gid1a* declined slower and continued for a few more days. When daily transpiration of each individual plant reached a minimum volume of 50 ml/day, irrigation was stopped completely. After three days of complete drought, we rehydrated the plants and plant recovery was monitored. Recovery was evaluated by the time required for each plant to return to full transpiration (the level of transpiration measured just before the beginning of the drought treatment; Negin and Moshelion, 2017). While *gid1a* plants were fully recovered within 3 days, M82 plants did not recover completely even after 10 days of irrigation (Fig. 3B).

**Fig. 3.**
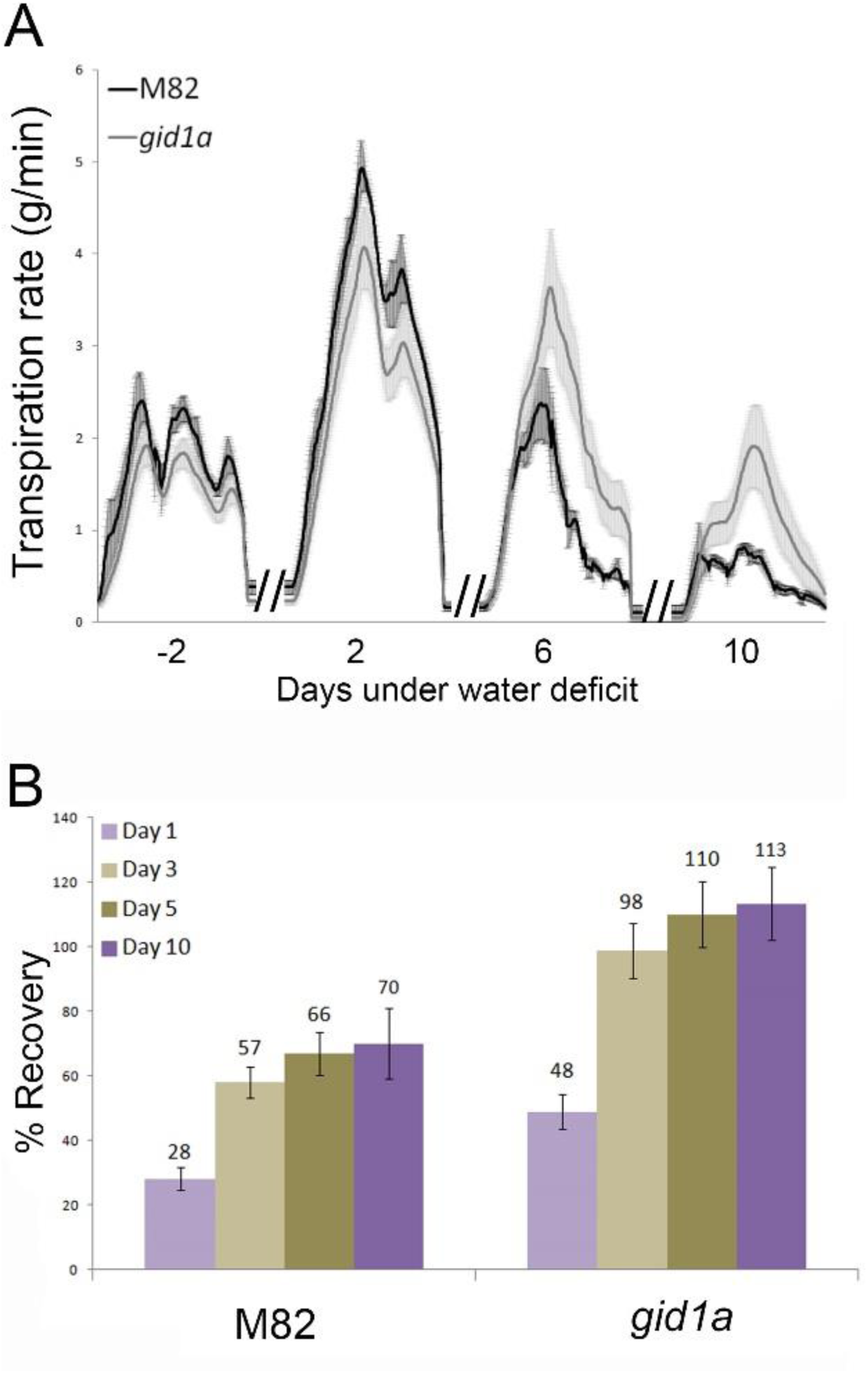
Loss of GID1a reduces transpiration under water deficit conditions. **A.** Transpiration rate in M82 and *gid1a* under water limited conditions. After two weeks of irrigation on the lysimeters, irrigation was gradually reduced (50% of the previous day transpiration, automatically controlled by the system for each plant separately) until it was completely stopped. Transpiration rate in selected representative days during the water deficit treatment are presented; 2 days before the beginning of the water-deficit treatment (−2) and 2, 6 and 10 days into the drought treatment. Values are means of eight plants ± SE. **B.** Recovery of M82 and *gid1a* from the drought treatment. Recovery was evaluated by the time required for each plant to return to the level of transpiration measured just before the beginning of the water-deficit treatment. Values are means of eight plants ± SE. Numbers above columns represent the percentage from maximum transpiration (see above).

### GID1 activity promote xylem vessel proliferation and hydraulic conductivity

We next explored if the loss of GA receptors affects additional factors that can be attributed to transpiration limitation. Previously we showed that mutation in GID1a inhibits root growth (Illouz-Eliaz *et al*., 2019). We therefore tested if the root system of the strongest double mutant *gid1a gid1b2* limits water uptake and water loss under water-deficit conditions. To eliminate the effect of the shoot, we grafted M82 scions on *gid1a gid1b2* and M82 rootstocks. Grafted plants were grown for two weeks under normal irrigation and then irrigation was stopped for dehydration. After four days, when plants started wilting, leaf RWC was measured. We did not find differences in the RWC between plants grafted on *gid1a gid1b2* or M82 rootstocks (Figure 4A). Moreover, all plants, regardless their rootstocks, wilted at the same time (Fig. 4B). These results suggest that *gid1* roots do not affect the rate of water loss under water-deficit conditions.

**Fig. 4.**
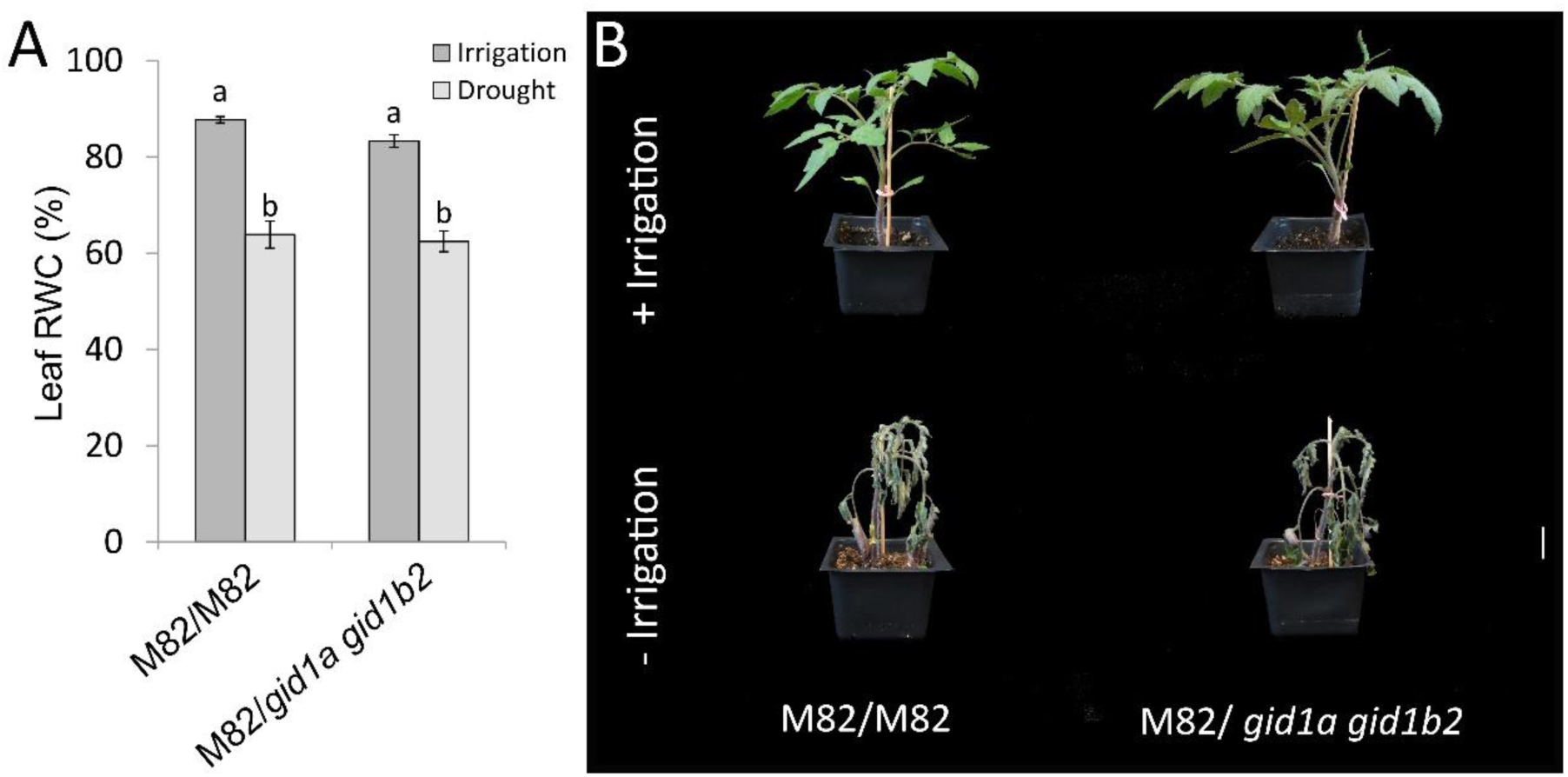
Grafting of M82 scions on M82 or *gid1a gid1b2* rootstocks. Grafted plants were grown for two weeks under normal irrigation (+irrigation) and then irrigation was stopped for dehydration (-irrigation). **A.** After four days, when plants started wilting, leaf RWC was measured. Values are mean of 6 plants ± SE. Small letters represent significant differences between the lines (Student’s t test, P<0.05). **B.** After seven days of the water-deficit treatment representative plants were photographed and are presented.

Since GA promotes secondary vascular development (Ragni *et al*, 2011; Dayan *et al*., 2012; Aloni, 2013), we examine if the loss of GID1s affects xylem development and hydraulic conductance. We first analyzed the xylem vessels in the leaf petioles of M82 and *gid1a* (leaf no. 4, top down). Microscopic analysis of total vessel area showed ca. 10% reduction in *gid1a* (Figure 5A). The reduced total xylem area was a results of reduced number of vessels (Figure 5B). We next evaluated how the reduced vessel number affects hydraulic conductance. To this end, we first calculated the specific theoretical hydraulic conductance of the xylem vessels in M82 and *gid1a*, using the Hagen-Poiseuille equation (Tyree and Ewers, 1991) and normalized it to the supported leaf area (Hochberg et al., 2015). The specific theoretical hydraulic conductivity of *gid1a* was ca. 23% lower than that of M82 (Fig. 5C). We then tested the actual hydraulic conductance, by measuring volumetric-flow rate in detached stem segments, taken from M82 and *gid1a* (Melcher *et al*., 2012). Hydraulic conductance of *gid1a* stems was ca. 20% lower than that of M82 (Fig. 5D). We also analyzed the stem vessel area and number in four-weeks-old M82 and *gid1a* plants. Total stem vessel area was 35% lower in *gid1a* due to 32% reduction in the number of xylem vessels (Fig. 5E and F, Supplementary Fig. S2A). The loss of GID1a did not affect xylem vessel expansion and the average area of individual xylem vessel in *gid1a* was similar to that in M82 (Supplementary Fig. S2B). To test if this is a general response to reduced GA activity, we analyzed xylem vessels and hydraulic conductance in transgenic plants overexpressing the stable DELLA protein proΔ17 (*35:proΔ17*, Nir *et al*., 2017). It should be noted that the inhibition of GA activity in *35:proΔ17* is much stronger than in *gid1a*. The number of vessels in *35:proΔ17* was 63% lower than in M82 (Supplementary Fig. S3A and B). In these transgenic plants, the reduced GA activity affected also vessel size and the average size of individual vessel was 26% lower than in M82 (Supplementary Fig. S3C). Total vessel area in *35:proΔ17* was ca. 70% lower than in M82 and hydraulic conductance (volumetric flow rate) ca. 80% lower (Supplementary Fig. S3D and E).

**Fig. 5.**
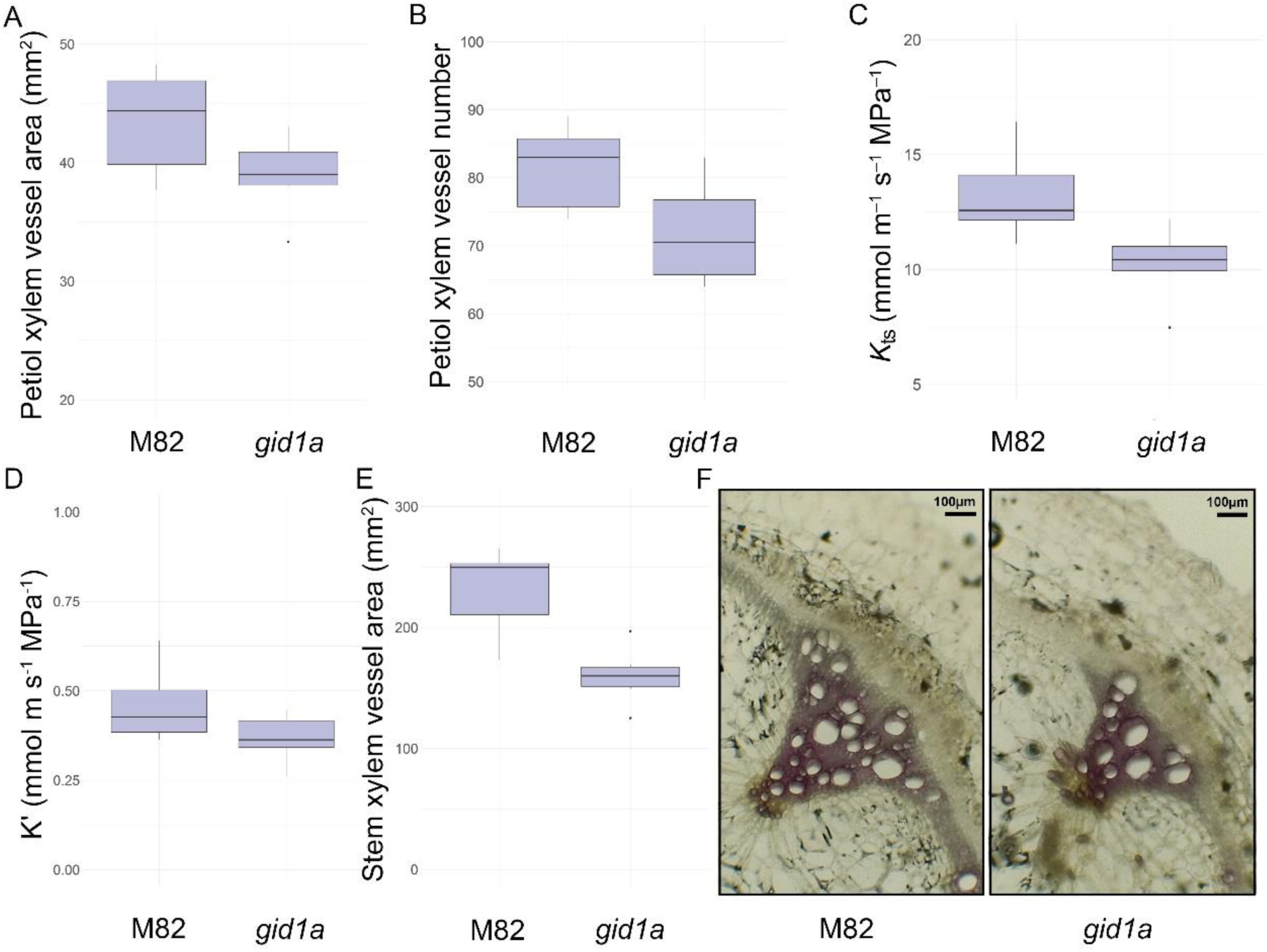
The loss of GID1a reduces xylem vessel proliferation and hydraulic conductance. **A.** Total xylem vessel area and **B**, xylem vessels number in the base of the leaf petioles of M82 and *gid1a* (leaf no. 4, top down). **C.** Theoretical specific hydraulic conductance (Kts) of the xylem vessels in M82 and *gid1a*. For calculation of Kt, the Hagen-Poiseuille equation (Tyree and Ewers, 1991) was used. To calculate Kts, Kt was normalized to the supported leaf area (Hochberg et al., 2015). **D.** Volumetric-flow rate was measured in detached stem segments, taken from four-weeks-old M82 and *gid1a* plants to calculate the actual hydraulic conductance (K’). **E.** Total xylem vessel area in M82 and *gid1a* stems. Data in **A**, **B**, **C D** and **E** (taken from 6 different plants) are graphically presented as whisker and box plots. **E.** Representative stem (as in **C**) cross-sections of M82 and *gid1a* stained with Wiesner stain. Scale bar = 100 μm.

To study further the effect of GA on xylem vessel development, we examined plant with even stronger reduction in GA activity. To this end, we have generated a CRISPR-Cas9 derived *sly1* mutant. SLY1 is the F-box that targets DELLA for degradation. Similar to Arabidopsis and rice, tomato has a single *SLY1*, (*SlSLY1*, Solyc04g078390, Liu *et al*., 2016). The mutations were analyzed by PCR and sequenced (Supplementary Fig. S4A). Homozygous mutant was obtained and the Cas9 construct was segregated out by back-crossing to M82. *sly1* has a single nucleotide insertion causing a frame shift prior to the LSL domain (Supplementary Fig. S4B), which is essential for the interaction with DELLA (Hirano et al., 2010). The homozygous *sly1* exhibited severe dwarfism and small dark-green leaves (Fig. 6A). *Sly1* exhibited insensitivity to exogenous treatment with 100μM GA_3_ (Supplementary Fig. S4C), suggesting strong inhibition of GA responses. To examine the effect of the reduced GA activity on xylem vessel development, we analyzed microscopically petioles of *sly1* and M82. Since *sly1* develops very slowly, we analyzed *sly1* and M82 petioles with similar diameter (the mutant leaves were much older). Figure 6B shows fewer and much smaller vessels in *sly1* compare to M82. These results suggest that reduced GA activity suppresses xylem vessel proliferation and expansion and these affect hydraulic conductance and probably limit transpiration.

**Fig. 6.**
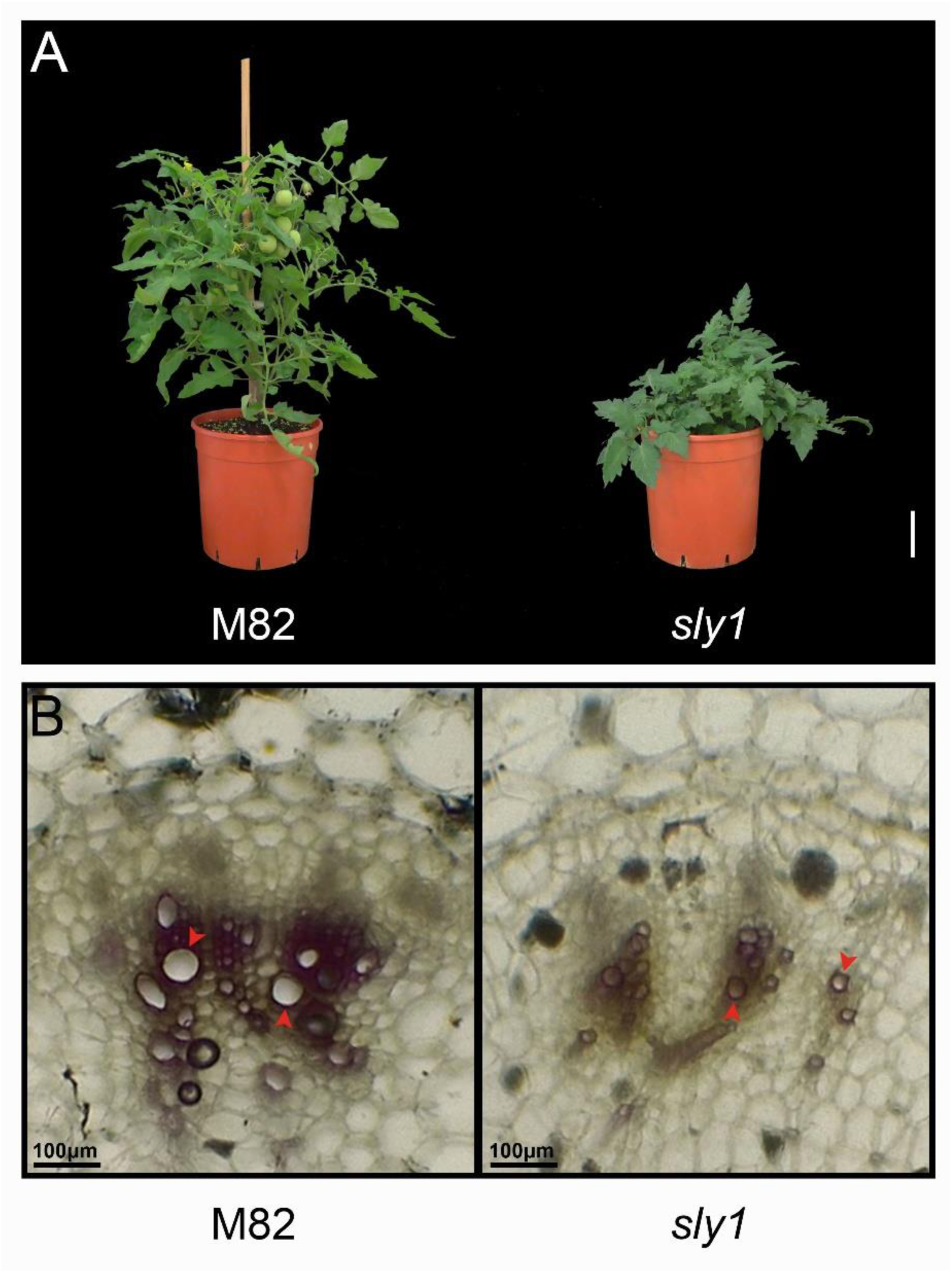
Loss of *SlSLY1* suppressed xylem vessel proliferation and expansion. **A.** Two-month-old M82 and representative CRISPR-Cas9 derived *sly1* mutant. Scale bar = 3 cm. **B.** Representative petiole cross-sections of M82 and *sly1* stained with Wiesner stain. *sly1* and M82 petioles with similar diameter (the mutant leaves were much older) were microscopically analyzed. Scale bar = 100 μm.

### *gid1*a in the field

We tested if the lower transpiration of the *gid1* mutant lines has advantage in the field, under drought conditions. M82 and all single mutant lines were planted in an open commercial field and the experiment was designed according to Gur and Zamir (2004). Fifteen plants from each line (M82, *gid1a, gid1b1* and *gid1b2*) were planted randomly and were irrigated normally throughout the experiment. Fifteen other plants of each line were irrigated normally for three weeks and then irrigation was stopped until harvesting (approximately three more months). It should be noted that during the drought treatment (May to August-Acre, Israel) no rain was recorded. Under normal irrigation regime, all single *gid1* mutant lines exhibited reduced growth compared to M82 (fresh weight, Fig. 7A). This loss of redundancy and semi-dwarfism of the *gid1*s under ambient conditions was reported by us before (Illouz-Eliaz *et al*., 2019). Despite this growth suppression, the single *gid1* mutant had similar fruit yield to M82 (green and red fruit, Fig. 7B). The drought treatment had stronger effect on M82 growth (as can be seen from the vegetative weight loss) compared to *gid1*s plants. However, M82 plants showed slightly higher vegetative fresh weight under drought conditions compared to all three *gid1* single mutants (Fig. 7A). The reduction in fruit yield under water-deficit conditions was similar in all lines (approximately 50% in M82 and all single *gid1* mutants, Fig. 7B). Lastly, we evaluated the parameter of harvest index (total yield per plant weight) for each line. *gid1a* showed significantly higher value of harvest index than all other lines, including M82 (Fig. 7C).

**Fig. 7.**
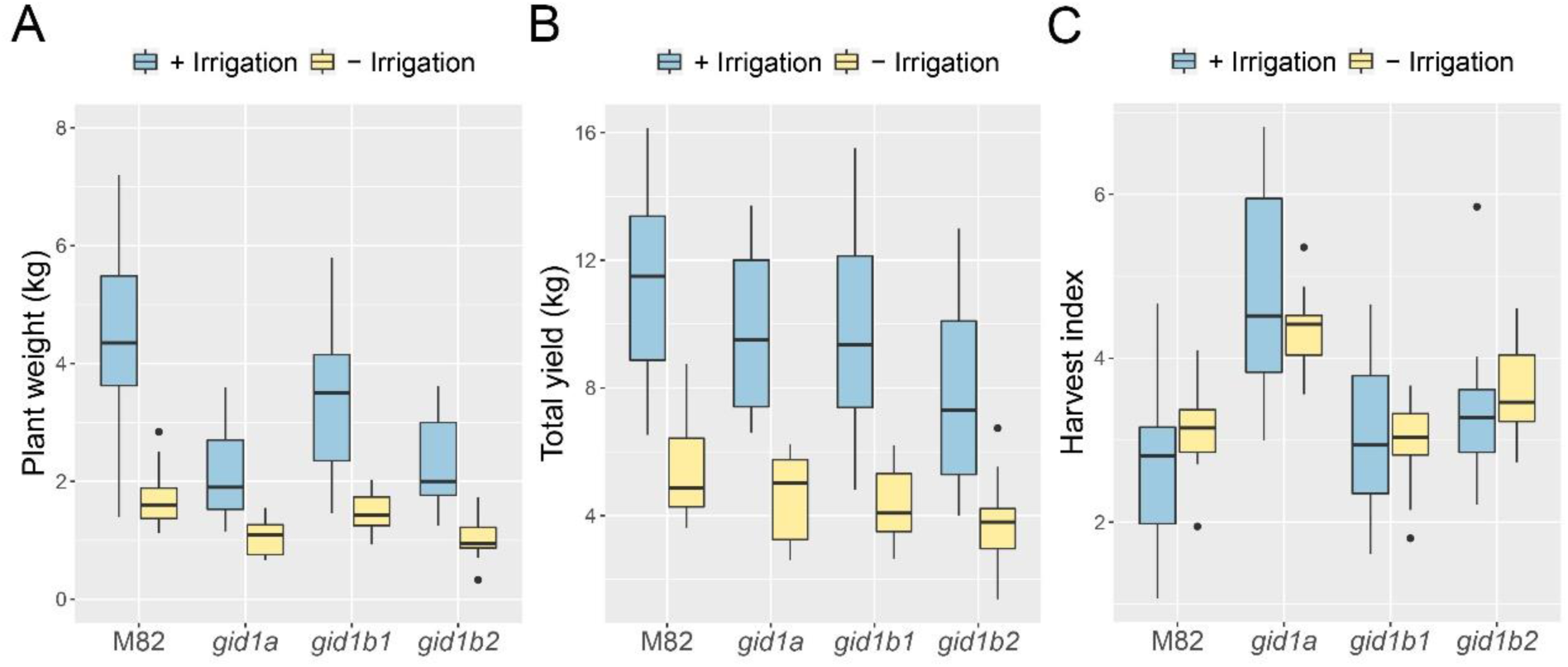
*gid1a* plants exhibit high harvest index in the field under irrigation and drought conditions. M82 and all single *gid1* mutant were planted in the field, Plants from each line were planted randomly and were irrigated normally throughout the experiment. Half of the plants of each line were irrigated normally for three weeks and then irrigation was stopped until harvesting (approximately three more months). **A.** Plant vegetative fresh weight (after removal of all fruits) at harvest. **B.** Total fruit yield (green and red fruits). **C.** Harvest index (total yield to vegetative fresh weight). Data in **A**, **B**, and **C** (taken from 15 different plants) are graphically presented as whisker and box plots.

## Discussion

Abiotic stresses, including drought, reduces GA levels and suppress plant growth (Colebrook *et al*., 2014). The reduced GA activity promotes tolerance to drought (Nir *et al*., 2017). Several possible mechanisms of how GA improve tolerance and/or drought avoidance were proposed, including reduced transpiration due to reduced plant size (Magome *et al*., 2008; Achard *et al*., 2006, Nir *et al*., 2014) and activation of various stress-related genes (Wang *et al*., 2008; Tuna *et al*., 2008). In tomato, reduced GA activity also promotes stomatal closure and reduces water loss under water-deficit conditions (Nir *et al*., 2014; Nir *et al*., 2017). It was suggested that accumulating DELLA (due to the reduced GA levels) promotes ABA responses in guard cells.

Low GA activity has a pleotropic effect on plant development. Since the three tomato GA receptors, GID1s have overlapping activities and high redundancy under normal growth conditions (Illouz-Eliaz *et al*., 2019), we examined here if mutation in *GID1* can improve water status under water-deficit conditions, without affecting growth and yield. Mutation in the most dominant GA receptor *GID1a* and its double mutants, *gid1a gid1b1* and *gid1a gid1b2* exhibited lower whole plant transpiration and reduced water loss under controlled water-deficit conditions (Fig. 2 and Illouz-Eliaz et al., 2019). The lower transpiration in *gid1a gid1b2* can be explained simply by the reduction in plant size. However, leaf area, stomatal density and stomatal index were not affected in *gid1a*. Thus, the reduced transpiration in this mutant probably resulted from reduced stomatal conductance.

Reduced hydraulic conductance of the xylem vessels leads to lower stomatal conductance and therefore, to reduced transpiration (Brodribb and Holbrook, 2003; Brodribb, 2009; Melcher *et al*., 2012). In Arabidopsis, GA promotes xylem-area expansion, due to secondary xylem differentiation (Ragni *et al*., 2011; Aloni, 2013). In tobacco stems, GA promotes cambial proliferation and secondary vascular development (Dayan *et al*., 2012). Here we show that mild suppression of GA activity in *gid1a* reduced xylem vessels number, which may explain the lower hydraulic conductance. In stronger GA lines (35S*:proΔ17* and *sly1* mutant) we found reduced number of vessels and reduced vessel size. The reduced number and size of vessels correlated well with the reduced GA activity (M82>*gid1a*>*35S:proΔ17>sly1)*. Thus, we suggest that decreased GA activity affect transpiration by multiple mechanisms; it reduces leaf area by inhibition of cell division and elongation, directly promotes stomatal closure by increasing ABA responses in guard cells (Nir *et al*., 2017) and indirectly, by reducing hydraulic conductance due to reduce xylem vessel number and size. While the mild attenuation of the GA signal in *gid1a* was not sufficient to inhibit stem elongation and leaf expansion, it was severe enough to suppress xylem vessel differentiation. This indicates that xylem vessel differentiation is extremely sensitive to changes in GA levels.

The lower transpiration found in the different GA mutants suggest that the improve performance of these lines under transient water deficit conditions is caused by drought avoidance (Kooyer, 2015). In the field however, the *gid1s* did not show advantage over the wild type M82 under drought conditions (fruit yield). Roots respond to water potential gradient and grow towards higher moisture content, a phenomenon called hydrotropism (Dietrich, 2018). In the field, the substantially larger root-zone increase the soil water reservoir, enabling roots to find new sources of moister. Thus, the plants sustained longer periods of water deficit conditions and were less dependent on the rate of transpiration.

While *gid1a* plants exhibited normal development in growth room, they were semi-dwarf in the field. This loss of redundancy under ambient conditions was demonstrated by us before; under extreme environmental conditions the activity of all three GID1s is required for robust growth (Illouz-Eliaz *et al*., 2019). Surprisingly, the decreased in growth of *gid1a* in the field did not affect fruit yield under both well-watered and water-deficit conditions and therefore, these plants showed the highest harvest index (fruit weight/plant fresh weight). Similarly, reduced GA activity suppresses growth but not yield in the ‘green revolution’ cereal varieties (Hedden, 2003; Harberd *et al*., 2009). These suggest that partial reduction in GA activity can restrict growth without affecting productivity. Harvest index is an important agronomic trait; it allows planting at higher density to obtain higher yield per unit area (Gifford and Evans, 1981). Thus, *gid1a* allele may be used to increase yield in cultivars with low harvest index. The potential of using the *gid1a* allele in breeding for higher yield requires further study.

## Supplementary data

Supplementary Table S1. List of primers and sgRNAs used in this study.

Supplementary Fig. S1. Stomatal density and index in M82 and *gid1a*.

Supplementary Fig. S2. Stem xylem vessel number and area in M82 and *gid1a*.

Supplementary Fig. S3. Xylem proliferation and expansion in *35:proΔ17*.

Supplementary Fig. S4. Analysis of the CRISPR-Cas9 derived *sly1* mutant.

## Acknowledgements

This research was supported by a research grants from the Israel Ministry of Agriculture and Rural Development (12-01-0014) and the Israel Ministry of Agriculture and Rural Development (Eugene Kandel Knowledge Center) as part of the “Root of the Matter”-The Root zone knowledge center for leveraging modern agriculture to D.W. We thank Dr. Uri Hochberg, Dr. Yotam Zait and Prof. Menachem Moshelion for helpful discussion. We also thank Shula Blum, Itamar Shenhar and Shai Turgeman for technical assistance.

## References

1. Achard P, Cheng H, De Grauwe L, Decat J, Schoutteten H, Moritz T, Van Der Straeten D, Peng J, Harberd NP. 2006. Integration of plant responses to environmentally activated phytohormonal signals. Science 311, 91–4.

2. Aloni R. 2013. Role of hormones in controlling vascular differentiation and the mechanism of lateral root initiation. Planta 238, 819–830.

3. Araujo WL, Nunes-Nesi A, Osorio S, Usadel B, Fuentes D, Nagy R, Balbo I, Lehman M, Studart-Witkowski C, Tohge T, Martinoia E, Jordana X, DaMatta FM, Fernie AR. 2011. Antisense inhibition of the iron-sulphur subunit of succinate dehydrogenase enhances photosynthesis and growth in tomato via an organic acid-mediated effect on stomatal aperture. The Plant Cell 23, 600–627.

4. Brodribb TJ, Holbrook NM. 2003. Stomatal closure during leaf dehydration, correlation with other leaf physiological traits. Plant Physiology 132, 2166–2173.

5. Brodribb TJ. 2009. Xylem hydraulic physiology: The functional backbone of terrestrial plant productivity. Plant Science 177, 245–251.

6. Colebrook EH, Thomas SG, Phillips AL, Hedden P. 2014. The role of gibberellin signalling in plant responses to abiotic stress. Journal of Experimental Biology 217, 67–75.

7. Cutler SR, Rodriguez PL, Finkelstein RR, Abrams SR. 2010. Abscisic acid: emergence of a core signaling network. Annual Review of Plant Biology 61, 651–679.

8. Davière J-M, Achard P. 2013. Gibberellin signaling in plants. Development 140, 1147–1151.

9. Dayan J, Voronin N, Gong F, Sun T-p, Hedden P, Fromm H, Alonia R. 2012. Leaf-induced gibberellin signaling is essential for internode elongation, cambial activity, and fiber differentiation in Tobacco stems. The Plant Cell 24, 66–79

10. Dietrich D. 2018. Hydrotropism: how roots search for water. Journal of Experimental Botany 69, 2759–2771.

11. Geisler M, Nadeau J, Sack FD. 2000. Oriented asymmetric divisions that generate the stomatal spacing pattern in Arabidopsis are disrupted by the too many mouths mutation. The Plant Cell 12, 2075–2086.

12. Gifford RM, Evans LT. 1981. Photosynthesis, carbon partitioning and yield. Annual Review of Plant Physiology 32, 485–509.

13. Griffiths J, Murase K, Rieu I, Zentella R, Zhang Z-L, Powers SJ, Gong F, Phillips AL, Hedden P, Sun T-p, Thomas SG. 2006. Genetic characterization and functional analysis of the GID1 gibberellin receptors in Arabidopsis. The Plant Cell 18, 3399–3414.

14. Gur A, Zamir D. (2004). Unused natural variation can lift yield barriers in plant breeding. PLoS Biology 2:e245. doi: 10.1371/journal.pbio.0020245

15. Halperin O, Gebremedhin A, Wallach R, Moshelion M. 2017. High-throughput physiological phenotyping and screening system for the characterization of plant–environment interactions. Plant Journal 89, 839– 850.

16. Harberd NP, Belfield E, Yasumura Y. 2009. The angiosperm gibberellin-GID1-DELLA growth regulatory mechanism: how an “inhibitor of an inhibitor” enables flexible response to fluctuating environments. The Plant Cell 21, 1328–1339.

17. Hauvermale AL, Ariizumi T, Steber CM. 2012. Gibberellin signaling: a theme and variations on DELLA repression. Plant Physiology 160, 83–92.

18. Hedden P. 2003. The genes of the Green Revolution. Trends in Genetics 19, 5–9.

19. Hirano K, Asano K, Tsuji H, Kawamura M, Mori H, Kitano H, Ueguchi-Tanaka M, Matsuoka M. 2010. Characterization of the molecular mechanism underlying gibberellin perception complex formation in rice. The Plant Cell 22, 2680–2696.

20. Hochberg U, Degu A, Gendler T, Fait A, Rachmilevitch S. 2015. The variability in the xylem architecture of grapevine petiole and its contribution to hydraulic differences. Functional Plant Biology 42, 357–365

21. Kooyer NJ. 2015. The evolution of drought escape and avoidance in natural herbaceous populations. Plant Science 234, 155–162

22. Illouz-Eliaz N, Ramon U, Shohat H, Blum S, Livne S, Mendelson D, Weiss D. 2019. Multiple gibberellin receptors contribute to phenotypic stability under changing environments. The Plant Cell 31, 1506–1519.

23. Liu Q, Guo X, Chen G, Zhu Z, Yin W, Hu Z. 2016. Silencing SlGID2, a putative F-box protein gene, generates a dwarf plant and dark-green leaves in tomato. Plant Physiology and Biochemistry 109, 491–501

24. Livne S, Lor VS, Nir I, Eliaz N, Aharoni A, Olszewski NE, Eshed Y, Weiss D. 2015. Uncovering DELLA-independent gibberellin responses by characterizing new tomato procera mutants. The Plant Cell 27, 1579–1594.

25. Locascio A, Blázquez MA, Alabadí D. 2013. Genomic analysis of DELLA protein activity. Plant and Cell Physiology 54,1229–1237.

26. Magome H, Yamaguchi S, Hanada A, Kamiya Y, Oda K. 2008. The DDF1 transcriptional activator upregulates expression of a gibberellin-deactivating gene, GA2ox7, under high-salinity stress in Arabidopsis. Plant Journal 56, 613–626.

27. McCormick S. 1991. Transformation of tomato with Agrobacterium tumefaciens. In Plant Tissue Culture Manual (ed. H. Linclsey), Kluwer Academic Publishers, Dordrecht, The Netherlands. pp 1–9.

28. Melcher PJ, Holbrook NM, Burns MJ, Zwieniecki MA, Cobb AR, Brodribb, TJ, Choat B, Sack L. 2012. Measurements of stem xylem hydraulic conductivity in the laboratory and field. Methods Ecol. Evol. 3: 685–694.

29. Mittler R, Blumwald E. 2010. Genetic engineering for modern agriculture: challenges and perspectives. Annual Review of Plant Biology 61, 443–462.

30. Negin B, Moshelion M. (2017). The advantages of functional phenotyping in pre-field screening for drought-tolerant crops. Functional Plant Biology 44, 107–118.

31. Nakajima M, Shimada A, Takashi Y, Kim YC, Park SH, Ueguchi-Tanaka M, Suzuki H, Katoh E, Iuchi S, Kobayashi M, Maeda T, Matsuoka M, Yamaguchi I. 2006. Identification and characterization of Arabidopsis gibberellin receptors. Plant Journal 46, 880–889.

32. Nir I, Moshelion M, Weiss D. 2014. The Arabidopsis GIBBERELLIN METHYL TRANSFERASE 1 suppresses gibberellin activity, reduces whole-plant transpiration and promotes drought tolerance in transgenic tomato. Plant Cell & Environment 37, 113–123.

33. Nir I, Shohat H, Panizel I, Olszewski N, Aharoni A, Weiss D. 2017. The Tomato DELLA protein PROCERA acts in guard cells to promote stomatal closure. The Plant Cell 29, 3186–3197.

34. Osakabe Y, Osakabe K, Shinozaki K, Tran L-S P. 2014. Response of plants to water stress. Frontiers in Plant Science 5, 86.

35. Omena-Garcia R, Martins A, Medeiros D, Vallarino J, Ribeiro D, Fernie A, Araujo W, Nunes-Nesi A. 2019. Growth and metabolic adjustments in response to gibberellin deficiency in drought stressed tomato plants. Environmental Experimental Botany 159, 95–107.

36. Pal S, Zhao J, Khan A, Yadav NS, Batushansky A, Barak S, Rewald B, Fait A, Lazarovitch N, Rachmilevitch S. 2016. Paclobutrazol induces tolerance in tomato to deficit irrigation through diversified effects on plant morphology, physiology and metabolism. Science Report 6, 39321.

37. Plaza-Wuthrich S, Blosch R, Rindisbacher A, Cannarozzi G, Tadele Z. 2016. Gibberellin deficiency confers both lodging and drought tolerance in small cereals. Frontiers in Plant Science 7, 643.

38. Pradhan Mitra P, Loqué D. 2014. Histochemical Staining of Arabidopsis thaliana Secondary Cell Wall Elements. Journal of Visualized Experiments 87, e51381.

39. Ragni L, Nieminen K, Pacheco-Villalobos D, Sibout R, Schwechheimer C, Hardtke CS 2011. Mobile gibberellin directly stimulates *Arabidopsis* hypocotyl xylem expansion. The Plant Cell 23, 1322–1336.

40. Sade N, Vinocur BJ, Diber A, Shatil A, Ronen G, Nissan H, Wallach R, Karchi H, Moshelion M. 2009. Improving plant stress tolerance and yield production: is the tonoplast aquaporin SlTIP2;2 a key to isohydric to anisohydric conversion? New Phytologist 181, 651–61.

41. Skirycz A, Inzé D. 2010. More from less: plant growth under limited water. Current Opinion in Biotechnology 21, 197–203.

42. Tuna AL, Kaya C, Dikilitas M, Higgs D. 2008. The combined effects of gibberellic acid and salinity on some antioxidant enzyme activities, plant growth parameters and nutritional status in maize plants. Environmental and Experimental Botany 62,1–9.

43. Tyree MT, Ewers FW. 1991. The hydraulic architecture of trees and other woody plants. New Phytologist 119, 345–360.

44. Ueguchi-Tanaka M, Ashikari M, Nakajima M, Itoh H, Katoh E, Kobayashi M, Chow T-y, Hsing Y-iC, Kitano H, Yamaguci I, Matsuoka M. 2005. *GIBBERELLIN INSENSITIVE DWARF1* encodes a soluble receptor for gibberellin. Nature 437, 693–698.

45. Wang C, Yang A, Yin H, Zhang J. 2008. Influence of water stress on endogenous hormone contents and cell damage of maize seedlings. Journal of Integrative Plant Biology 50, 427–434.

46. Wang R-S, Pandey S, Li S, Gookin TE, Zhao Z, Albert R, Assmann SM. 2011. Common and unique elements of the ABA-regulated transcriptome of Arabidopsis guard cells. BMC Genomics 12, 216.

47. Weber E, Engler C, Gruetzner R, Werner S, Marillonnet S. 2011. A modular cloning system for standardized assembly of multigene constructs. PLoS One. 6, e16765.

