## Supplemental Figure S1 for "Mutations in the tomato gibberellin receptors suppresses xylem proliferation and reduces water loss under water-deficit conditions"

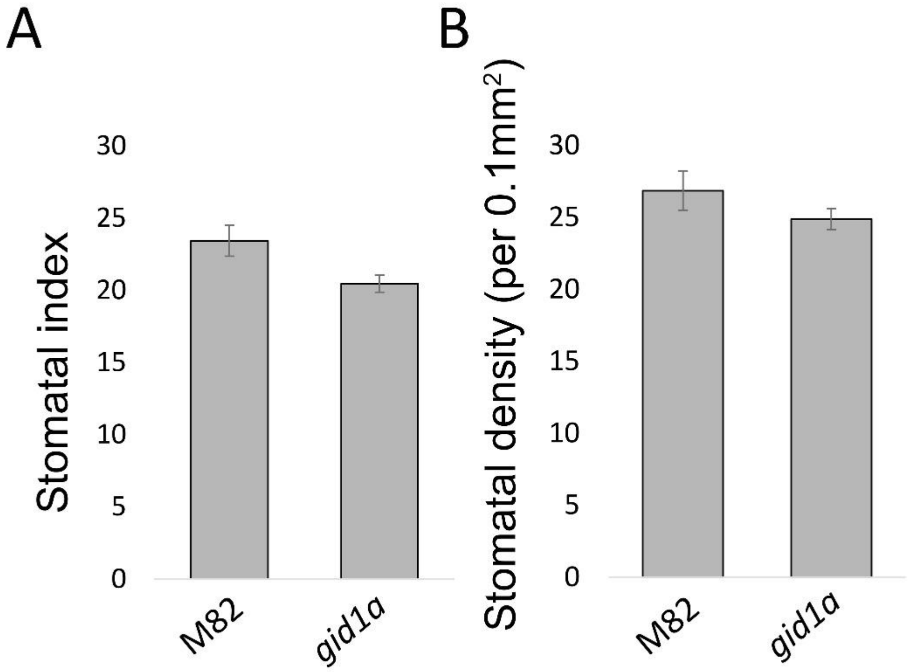


**Fig. S1.** The loss of GID1a has no effect on stomatal density or index. **A.** Stomatal density (per 0.1 $mm^2$). **B**. Stomatal index (number of stomata to epidermal cells). Values are means of five replicates (5 different plants) ± SE.
