## Supplemental Figure S2 for "Mutations in the tomato gibberellin receptors suppresses xylem proliferation and reduces water loss under water-deficit conditions"

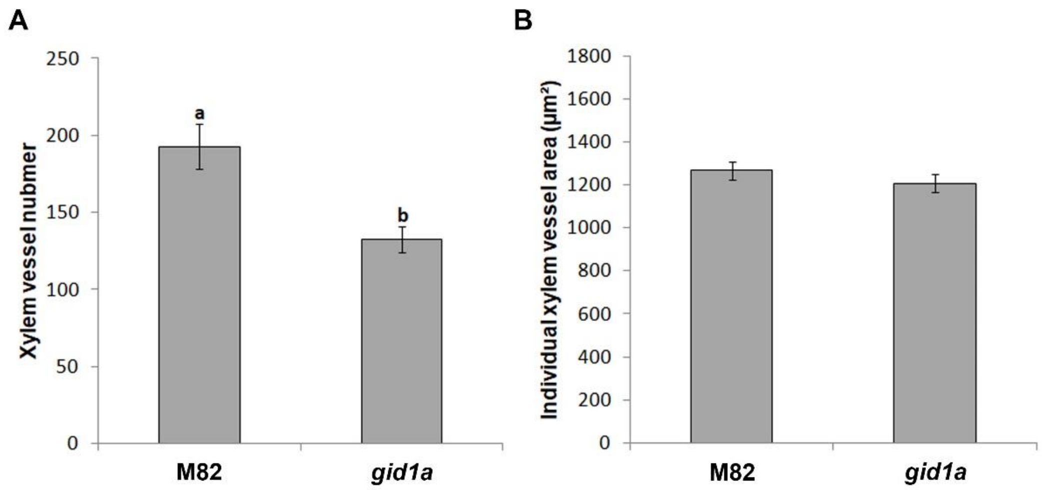


**Fig. S2. A.** Xylem vessel number in M82 and *gid1a* stems. **B.** Area mean of individual xylem vessel in M82 and *gid1a* stems. Values in **A** and **B** are means of 6 plants ± SE.
