## Supplemental Figure S3 for "Mutations in the tomato gibberellin receptors suppresses xylem proliferation and reduces water loss under water-deficit conditions"

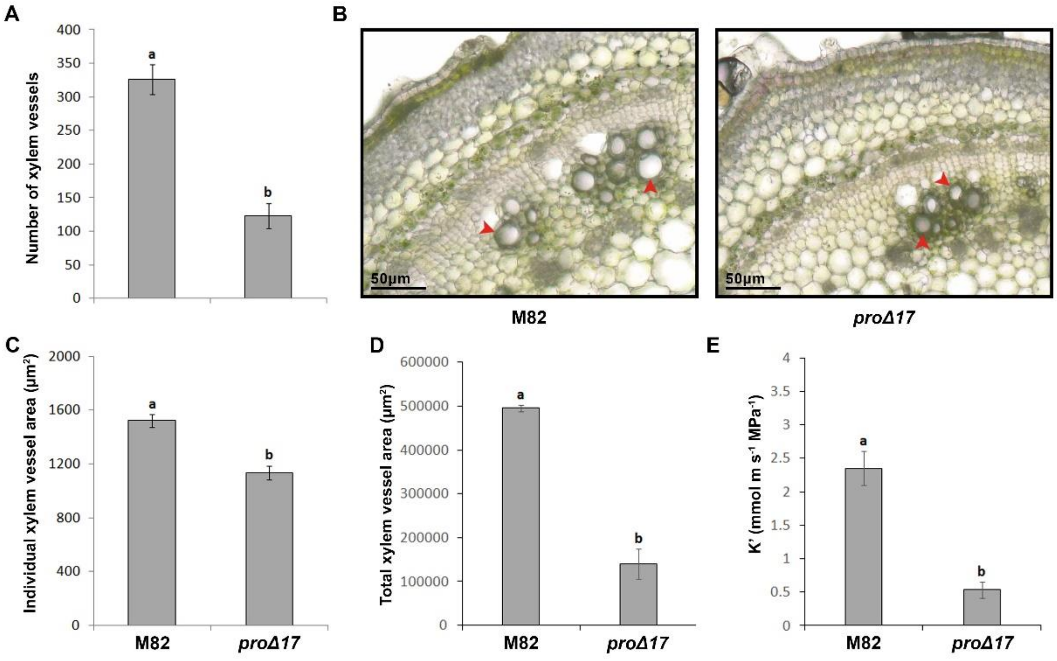


**Fig. S3. A.** Xylem vessel number in the stems of five-weeks-old M82 and *35:proΔ17* plants. **B.** Representative stem cross-sections of M82 and *35:proΔ17*. Scale bar = 50 µm. **C**. Area mean of individual xylem vessel in M82 and *35:proΔ17* stems**. D.** Total xylem vessel area in M82 and *35:proΔ17* stems. **E.** Hydraulic conductance measured in detached stem segments, taken from five-weeks-old M82 and *35:proΔ17*. Values in **A**, **C**, **D** and **E** are means of 6 plants ± SE. Small letters represent significant differences between the lines (Student's t test, $P<0.05$).
