## Supplemental Figure S4 for "Mutations in the tomato gibberellin receptors suppresses xylem proliferation and reduces water loss under water-deficit conditions"

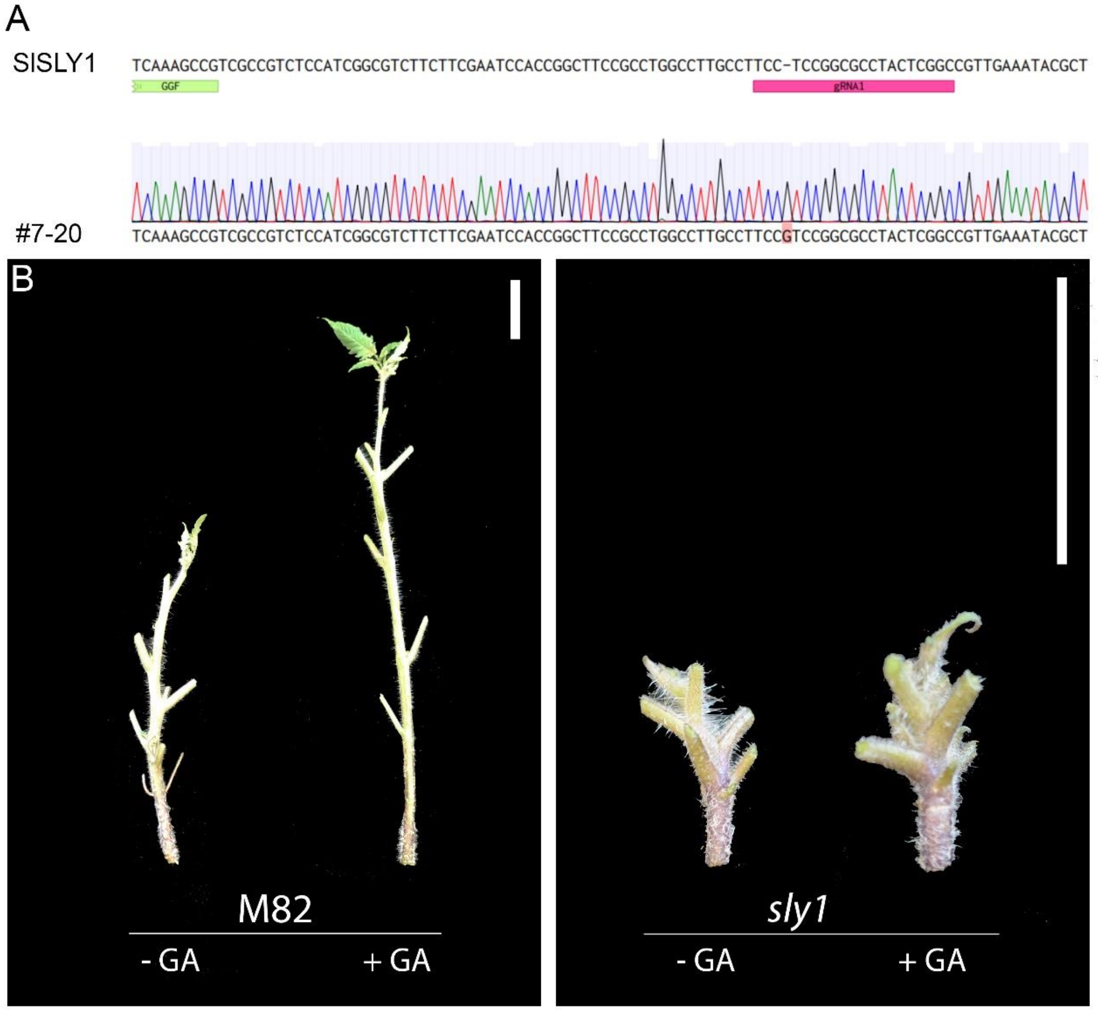


**Fig. S4.** Molecular and phenotypic characterization of *sly1*. Sequence of *SlSLY1* mutant alleles. **A.** Sequences of SLY1 wild-type (upper) and *sly1* mutant allele #7-20 with one nucleotide deletion (G, marked pink). Pink bar represents RNA guide sequence. **B.** Effect of 100 μM GA3 treatment on the elongation of M82 and *sly1*. Leaves were removed to show internode elongation. Scale bar = 2 cm.
