## Supplemental Table 1 for "Mutations in the tomato gibberellin receptors suppresses xylem proliferation and reduces water loss under water-deficit conditions"

**Supplementary data**

**Table S1.** Primers used in this study.

| Gene | Used for | Sequence (5'-3') |
|---|---|---|
| *GID1a* | qRT-PCR | Forward- TCTTGTTGTTGTCGCAGGTT<br>Reverse- CCCTATTGTTGCCTTCTCCA |
| *GID1b1* | qRT-PCR | Forward- GGCTGCTCTTCAATGGGTAA<br>Reverse- TAAAACCTCGACGCCTGATT |
| *GID1b2* | qRT-PCR | Forward- TGAGGCTGATTGGGGTAAAA<br>Reverse- TAGGCGGCGACAAAATGTAT |
| *SlACTIN* | qRT-PCR | Forward- GTCCTCTTCCAGCCATCCAT<br>Reverse- ACCACTGAGCACAATGTTACCG |
| *SlSLY1* | CRISPR/ Cas9 | Guide 1- GCCGAGTAGGCGCCGGAGGA<br>Guide 2- GGTTATAGAGTCGCCGGATG |
| *SlSLY1* | Cloning to pBridge | Forward-<br>ATGAAGCGGCAATTCGAC<br>Reverse-<br>TTATTTATTAGCTTTGAAATTCATCTTCTCGTAG |
